# Mammarenavirus-Induced Remodeling of the Cellular Lipid Landscape Reveals Sphingolipid Metabolism as a Novel Target for Antiviral Intervention

**DOI:** 10.64898/2026.07.02.736094

**Authors:** Patricia Mingo-Casas, Haydar Witwit, Mireia Casasampere, Ana-Belén Blázquez, Beatrice Cubitt, Miguel A. Martín-Acebes, Juan Carlos de la Torre

**Affiliations:** Department of Biotechnology, Instituto Nacional de Investigación y Tecnología Agraria y Alimentaria, Consejo Superior de Investigaciones Científicas (INIA-CSIC), 28040, Madrid, Spain; Universidad Autónoma de Madrid (UAM, Escuela de Doctorado), Madrid, Spain; Department of Immunology and Microbiology, The Scripps Research Institute, La Jolla, CA 92037, USA; Department of Biological Chemistry, Institute for Advanced Chemistry of Catalonia (IQAC), CSIC, Barcelona, Spain; CIBER of Hepatic and Digestive Diseases (CIBEREHD), Instituto de Salud Carlos III (ISCIII), Madrid, Spain

**Keywords:** Mammarenavirus, LCMV, lipid, sphingolipid, sphingomyelinase, host-directed antiviral

## Abstract

Several mammarenaviruses (MaAv) cause severe and often life-threatening disease in humans and represent major public health threats in their endemic regions. Lassa (LASV) and Junin (JUNV) MaAv, endemic to Western Africa and the Argentine Pampas, respectively, are etiologic agents of viral hemorrhagic fevers associated with high morbidity and mortality. In addition, the globally distributed MaAv lymphocytic choriomeningitis virus (LCMV) is an underrecognized human pathogen capable of causing severe congenital disease and fatal infections in immunocompromised individuals. Despite their public health importance, no FDA-approved vaccines or virus-specific antiviral therapies exist to prevent and treat human MaAv infections. Current treatment relies on the off-label use of ribavirin whose therapeutic efficacy remains controversial. These findings underscore the urgent need to develop effective antiviral strategies against human pathogenic MaAv. Here, we investigated the impact of LCMV infection on host lipid metabolism using an integrated transcriptomic and lipidomic approach. Our data reveal extensive time-dependent remodeling of the cellular lipid landscape, with particularly prominent alterations in sphingolipid and fatty acid metabolic pathways. Functional interrogation of these pathways using pharmacological inhibitors identified acetyl-CoA carboxylase (ACC) and neutral sphingomyelinase 2 (nSMase2) as host factors contributing to efficient viral replication. Notably, inhibition of nSMase2 reduced infectious virus production by 2 logs of infectious virus. Our findings showed that LCMV reprograms host lipid metabolism to facilitate infection and identified sphingolipid turnover as a promising target for host-directed antiviral strategies against MaAv infections.

## 1. INTRODUCTION

Mammarenaviruses (MaAv) cause chronic infections in their rodent reservoirs worldwide with human infections occurring primarily through mucosal exposure to aerosolized infectious material or direct contact with contaminated materials (Radoshitzky SRB & de la Torre, 2022). Several MaAv cause hemorrhagic fever (HF) diseases in humans and represent important public health challenges in their endemic regions. The Old World (OW) MaAv Lassa virus (LASV), endemic to Western Africa, is estimated to infect over 500,000 individuals annually (Basinski et al., 2021). LASV ranks very high among zoonotic viruses with the potential for spillover and spread in humans (Grange et al., 2021), and its endemic regions are expanding (Sogoba et al., 2012; Klitting et al., 2022). The inclusion of LF on the WHO R&D Blueprint priority list further underscores the urgent need for the development of antivirals against LASV (Sweileh, 2017). Likewise, the New World (NW) MaAv Junin (JUNV), the causative agent of Argentine HF (AHF), and several other NW MaAv cause severe diseases throughout South America (Grant, A. et al., 2012; Lendino et al., 2024). In addition, evidence indicates that the worldwide-distributed prototypic MaAv lymphocytic choriomeningitis virus (LCMV) is an underrecognized human pathogen of clinical significance, particularly in congenital infections, and LCMV can cause severe disease in immunocompromised individuals (Bonthius, 2009; Palacios et al., 2008).

Despite their impact on human health, there are currently no FDA-approved MaAv vaccines or antivirals. Current treatment for LASV infections is limited to the off-label use of ribavirin (RBV) whose efficacy remains controversial (Salam et al., 2021). Several direct-acting antivirals (DAAs) including the polymerase inhibitors favipiravir (Mendenhall et al., 2011; Rosenke et al., 2018; Safronetz et al., 2015), and 4’-fluorouridine (4’-FIU) (Lieber & Plemper, 2022), as well as entry inhibitors LHF-535 (Cashman et al., 2022) and ARN-75039 (Gowen et al., 2021), and antibody-based therapies (Cross et al., 2019) are under development and have shown promising results in preclinical animal models of LF and AHF. However, clinical development of these DAAs may be complicated by toxicity and resistance emergence. Likewise, implementation of antibody-based therapies may be limited by high cost, dependence on reliable cold chain distribution, and the need for parenteral delivery, which may restrict their use to specialized treatment centers. Therefore, the development of broad-spectrum antivirals to counteract human pathogenic MaAv represents a significant unmet biomedical need.

Host-directed antivirals (HDAs) represent a promising yet underexplored approach in antiviral research (Kumar et al., 2020; Zumla et al., 2016). In contrast to conventional DAAs, which directly target viral proteins and functions, HDAs disrupt host factors and cellular processes hijacked by viruses to support their replication and pathogenesis. Since members of a virus family share host dependencies, HDAs have the potential to act as broad-spectrum antivirals. In addition, HDAs impose a higher genetic barrier to the emergence of drug-resistant viral variants, which often compromise antiviral therapy (Domingo et al., 2006; Kaufmann et al., 2018; Kai et al., 2017). Lipid metabolism represents an attractive target, as many viruses depend on extensive remodeling of host lipid pathways for replication, assembly, and immune evasion (Lai et al., 2022; Husby & Stahelin, 2021). The potential of lipids as antiviral targets is supported by their proven druggability and established clinical safety for the treatment of other human diseases (Blázquez et al., 2025). Lipid metabolism has been implicated in the regulation of T cell responses to LCMV (Kudek et al., 2023; Kazane et al., 2024; Harabuchi et al., 2022). Additionally, sphingomyelin has been shown to play a proviral role in LCMV-infected cells (Han, S. et al., 2024), and during the preparation of this manuscript, MaAv dependence on fatty acid synthesis was independently described in (Noble et al., 2026). Defining the lipid pathways required for optimal MaAv multiplication can identify novel targets for the development of HDAs against human pathogenic MaAv. In this study, we combined transcriptomic and lipidomic approaches to characterize the impact of LCMV infection on host lipid metabolism and identified acetyl-CoA carboxylase (ACC) and neutral sphingomyelinase 2 (nSMase2) as druggable targets for the development of HDAs against MaAv. Future validation studies in human primary cells and animal models of MaAv infection will be required to assess the translational potential of the findings here reported.

## 2. MATERIALS AND METHODS

### 2.1. Cells, viruses and infections

Vero E6 (*Chlorocebus aethiops*; ATCC CRL-1586) and A549 (*Homo sapiens;* ATCC CCL-185) cell lines were maintained in Dulbecco’s modified Eagle medium (DMEM) (ThermoFisher Scientific) containing 10% heat-inactivated fetal bovine serum (FBS), 2 mM L-glutamine, 100 μg/mL streptomycin, and 100 U/mL penicillin. FluoroBrite DMEM was purchased from Life Technologies. Armstrong strain (clone 5) of LCMV (Lee et al., 2002) (GenBank: NC-077806.1 and NC-077807.1) and recombinant rLCMV/GFP-P2A-NP (Witwit et al., 2024) have been described. Virus infections were performed on cell monolayers at a multiplicity of infection (MOI) of 0.05. After 1 hour adsorption at 37 °C, then viral inoculum was removed and cells were fed with fresh medium. Cells were maintained 24 or 48 hours at 37°C and 5% CO_2_.

### 2.2. Chemical compounds

PF-05175157, CP-640186, Dimethoxy-4-(5-Phenyl-4-Thiophen-2-yl-1H-Imidazol-2-yl)-Phenol (DPTIP) and Cambinol were purchased from MedChemExpress, resuspended in 1 ml of DMSO (10 mM) stocks and stored at -20°C until its use. Ribavirin (RBV) was purchased from Sigma-Aldrich.

### 2.3. Antiviral activity and cytotoxicity assessment

Vero E6 cells were seeded at 4 × 10^4^ cells/well into a 96-well plate and infected with rLCMV/GFP (MOI 0.05 FFU/cell). After a 90 min adsorption, the virus inoculum was aspirated off and serial dilutions (2-fold) of inhibitors were added to the cells. We used treatment (100 µM) with RBV as a validated inhibitor of mammarenavirus multiplication (Ofodile et al., 2025). Cell viability was determined by using the CellTiter 96 AQueous One Solution Reagent (Promega) (Witwit et al., 2024). The half-maximal cytotoxic concentrations (CC_50_) and half-maximal effective concentrations (EC_50_) were determined using GraphPad Prism, v10 (Prism10).

### 2.4. Immunofluorescence imaging

Hoechst (1:1000 in FluoroBrite DMEM medium) was used to stain nuclear DNA (Witwit et al., 2024). Immunofluorescence images (4X magnification) were taken using Keyence BZ-X710 fluorescence microscope.

### 2.5. RNA-seq and transcriptomic analysis

Total cellular RNA was extracted from A549 cells infected with rLCMV/GFP (MOI 0.05 FFU/cell) at 24 at 48 hours post infection (hpi) using the TRI reagent (TR 118) (Molecular Research Center). RNA was resuspended in sodium citrate pH 6.4 and stored at −80 °C until its use. cDNA libraries were generated from RNA samples by using the Zymo-Seq RiboFree Total RNA Library Kit (Zymo Research) and sequencing done using the Illumina NovaSeq X platform. Differentially expressed genes (DEGs) were selected by using DESeq2 (Love et al., 2014) based on |log2 fold change infected/uninfected|> 2 and adjusted P-value < 0.05 corrected by false discovery rate (FDR) values (Mingo-Casas, Blázquez et al., 2023). Principal component analysis (PCA) plots were performed with SRplot (Tang et al., 2023). Graphs displaying DEG intersection between lipid metabolism gene set R-HSA-556833.9, retrieved from Reactome knowledge database (Milacic et al., 2024), and infected cells datasets were created with Venn diagram generator online tool (https://bioinformatics.psb.ugent.be/webtools/Venn/). Functional enrichment and protein–protein networks were created using STRING (Szklarczyk et al., 2023) and analyzed in Cytoscape 3.10.3 (Shannon et al., 2003) using stringApp (Doncheva et al., 2019). Gene ontology (GO) enrichment for Biological Process (GO:BP) was performed using STRING online tool (Szklarczyk et al., 2023) and bubble plot showing gene count and –log_10_ (qvalue) was created with SRplot. GO:BP enriched terms were filtered with REVIGO (Supek et al., 2011) to identify non-redundant terms. Transcription factor enrichment for multiple gene lists and heatmap clustering was performed using METASCAPE (Zhou et al., 2019) and TRRUST database (Han, H. et al., 2018).

### 2.6. Lipidomics

A549 cells infected with LCMV Arm or uninfected controls were collected at 24 and 48 hpi. Lipidomic analyses were performed using a cell suspension at a concentration of 2 × 10⁶ cells/mL and following the protocol described in (Casasampere et al., 2026). Fold change in lipid levels between control and infected samples was calculated as log_2_ (treated/control) and data were analyzed with Metaboanalyst 6.0 web-based platform (Pang et al., 2024). Lipid annotations were performed according to (Liebisch et al., 2020) as follows: Sphingolipids (Cer, ceramide; dhCer, dihydroceramide; SM, sphingomyelin; dhSM, dihydrosphingomyelin; hexCer, hexosylceramide; lacCer, lactosylceramide; CTH, ceramide trihexoside; GM1 and GM2, gangliosides), glycerophospholipids (PC, phosphatidylcholine; PCO, ether-linked phosphatidylcholine; LPC, lysophosphatidylcholine; PE, phosphatidylethanolamine;) and sterols (CHOL, cholesterol).

### 2.7. Viral growth kinetics

A549 cells were infected at MOI 0.05 and treated with indicated compounds at specified concentrations. At designated times post-infection, cell culture supernatants (CCS) were collected, cells were fixed, and viral titers determined by focus-forming assay (FFA) as previously described (Witwit et al., 2026).

### 2.8. Virus titration

Virus titers were determined by focus-forming assay (FFA) as described (Witwit et al., 2024; Battegay et al., 1991). Titers of LCMV-ARM LCMV were determined by FFA using the rat monoclonal antibody VL4 to NP (Bio C Cell) and Alexa Fluor 488 anti-rat as secondary antibody.

### 2.9. Data analysis

For statistical comparisons, two-way analysis of the variance (ANOVA) with Sidak multiple comparison tests approach or ordinary ANOVA with Sidak multiple comparison were performed with Prism 10.0 for Windows (GraphPad Software, Inc.). Statistically significant differences are denoted in the figures (*, P < 0.05; **, P < 0.01; ***, P < 0.001; ****, P < 0.0001).

## 3. RESULTS

### 3.1. Effect of LCMV infection on host cell lipid metabolism pathways

To assess the impact of LCMV infection on the host cell lipid metabolism we performed RNA-seq analysis of LCMV-and mock-infected control A549 cells. We infected (MOI = 0.05) A549 cells with a recombinant LCMV expressing GFP, rLCMV/GFP-P2A-NP (Kim et al., 2019; Miranda et al., 2018), and monitored infection progression based on virus directed GFP expression (Fig.1A). At 24 and 48-hours post-infection (hpi) we determined titers of infectious virus in cell culture supernatant (CCS) samples by focus forming unit assay (FFA) (Fig. 1B) and stained cells with Hoechst in FluoroBrite DMEM (Fig. 1B). After images acquisition, we isolated total cellular RNA from each sample and performed bulk RNA sequencing. Principal components analysis (PCA) demonstrated clear separation between the gene expression profiles of LCMV-infected and mock-infected cells (Fig. 1C) with PC1 and PC2 accounting for 58% of variance (PC1=38.2%, PC2=19.9%); remaining variance reflects biological heterogeneity in infection efficiency. The number of differentially expressed genes (DEGs) increased between LCMV and mock-infected control cells over the course of the infection (Fig. 1D), with 230 DEGs detected at 24 hpi and up to 1,310 at 48 hpi. We performed functional enrichment of DEGs identified at 48 hpi against the Gene Ontology Biological Process (GO:BP) followed by subsequent filtering of redundant terms. The top ten terms were related to defense response to pathogens, responses to stimuli, and to stress, and immune responses (Fig. 1E). Additionally, we observed enrichment of a variety of terms related to different aspects of lipid metabolism such as fatty acid, arachidonic and eicosanoid metabolism, as well as lipid biosynthesis, regulation and storage (Fig. 1F). We further investigated the specific DEGs related to lipid metabolism by integrating our data with Reactome pathway annotations (Fig. 2A), which resulted in the identification of 40 lipid metabolism-associated DEGs, with 6 of them observed at 24 and 48 hpi, whereas 34 were only detected at 48 hpi. Protein-protein interaction (PPI) network for the 40 lipid metabolism-associated DEGs further highlighted a coordinated regulation of pathways involved in fatty acid and sphingolipid metabolism suggesting extensive rewiring of host lipid networks during LCMV infection (Fig. 2B).

**Figure 1.**
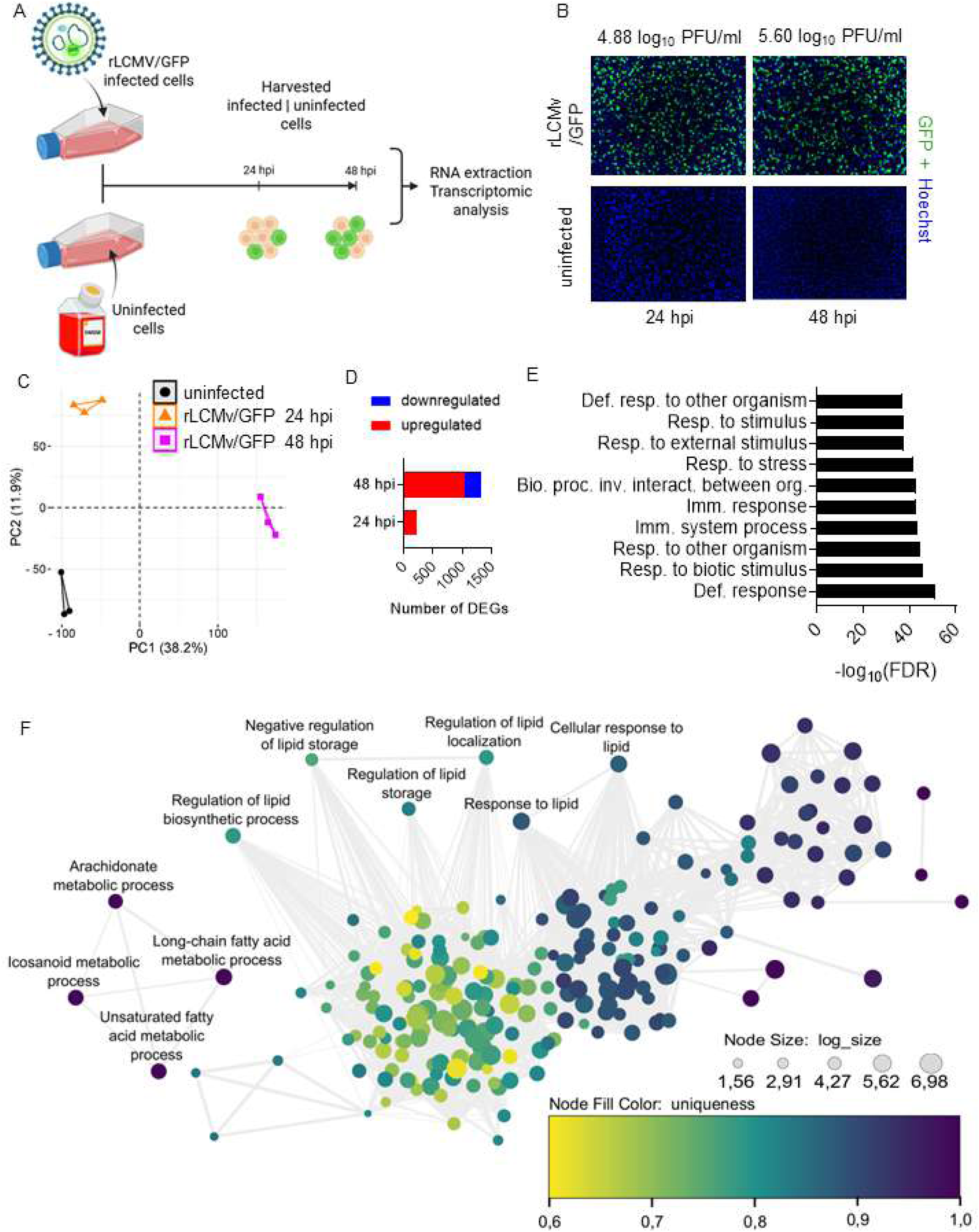
Transcriptomic reprogramming upon LCMV infection rewires pathways related to lipid metabolism. **A.** Experimental design and sample collection. A549 cells were infected with rLCMV/GFP (MOI = 0.05) or mock infected (DMEM). Figure created with Biorender. **B.** Representative immunofluorescence images of rLCMV/GFP and mock-infected control cells at 24 and 48 hpi. Hoechst was used to stain nuclear DNA. Images were taken at 4× magnification using a Keyence BZ-X710 fluorescence microscope. **C.** Principal component analysis (PCA) plot showing gene expression profiles in A549 cells at 24 and 48 hpi. **D.** Total number of differentially expressed genes (DEGs) identified using criteria |FC|> = 2 and FDR-corrected P < 0.05. Upregulated (red) and downregulated (blue) genes correspond to FC > 2 and FC < 2, respectively, relative to mock-infected control cells. **E.** Top 10 enriched GO:BP at 48 hpi GO:0006952 (Defense response); GO:0009607 (Response to biotic stimulus); GO:0051707 (Response to other organism); GO:0002376 (Immune system process); GO:0006955 (Immune response); GO:0044419 (Biological process involved in interspecies interaction between organisms); GO:0006950 (Response to stress); GO:0009605 (Response to external stimulus); GO:0050896 (Response to stimulus); GO:0098542 (Defense response to other organism).**F.** Interaction network of enriched lipid metabolism-related GO:BP terms identified among all DEGs at 48 hpi. The network was filtered to display the largest subnetwork using REVIGO software. Nodes are colored based on uniqueness (calculated as 1 – (average semantic similarity of a term to all other terms)) and node size represents log size (Log10; number of annotations for GO term ID in selected species in the EBI GOA database).

**Figure 2.**
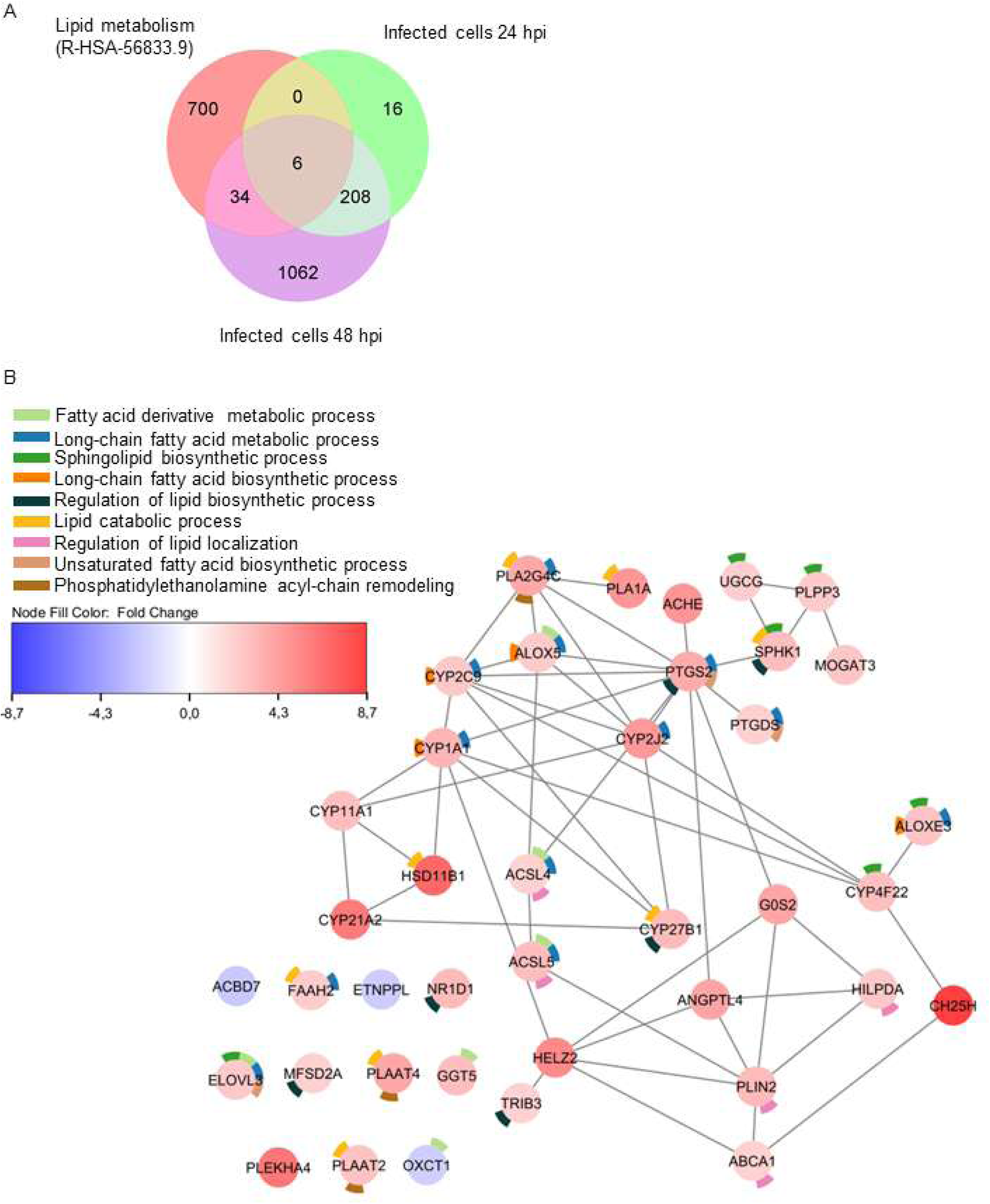
Temporal changes in lipid metabolism during LCMV infection. **A.** Venn diagram displaying the overlap between DEGs identified at 24 and 48 hpi in A549 cells and “Lipid Metabolism” Reactome gene set (R-HSA-556833.9). **B.** PPI network generated for the 40 lipid metabolism-related DEGs identified at 24 and 48 hpi. Nodes are colored based on fold change and gene symbol are showed inside each node. Donut plot segments indicate selected over-represented GO:BP terms identified by STRING enrichment analysis.

### 3.2. Effect of LCMV infection on the cellular lipidome

To determine whether transcriptional changes translated into alterations in lipid composition, we performed lipidomic profiling of LCMV and mock-infected cells at 24 and 48 hpi. For this we infected (MOI = 0.05) A594 cells with LCMV, or mock-infected, and at 24 and 48 hpi we collected CCSs and cells (Fig. 3A). Infection progression was monitored by titration of infectious progeny in CCSs (Fig. 3B). Cellular lipids were extracted, identified and quantified by LC-MS/ToF. This analysis covered four lipid classes spanning sphingolipids, glycerophospholipids, storage lipids (glycerolipids) and sterol lipids. We identified a total of 101 molecular species, covering 16 lipid subclasses (Fig. S1). At the lipid subclass level (Fig. 3C), we detected significant increases in levels of ceramides (Cer), sphingomyelin (SM), dihydrosphingomyelin (dhSM), lactosylceramides (lacCer), ganglioside (GM2) and ceramide trihexoside (CTH) in LCMV-infected cells at 48 hpi. Notably, among the other lipid classes analyzed we observed a significant increase only in lysophosphatidylcholine (LPC) at 24 hpi. The most notable changes were an increase in LPC and PE species at 24 hpi, and an increase in multiple sphingolipids (GM1, laCer, dhCer, GM2, Cer, hexCer, CTH and GM2) and LPC and PC species at 48 hpi (Fig. 3D). Taken together, these results support changes in the content of selective remodeling of cellular lipidome in LCMV infected cells, pointing to sphingolipids as the predominant lipid class altered during LCMV infection. Concurrent SM and Cer accumulation indicates complex sphingolipid remodeling controlled post-translationally, independent of transcriptional changes.

**Figure 3.**
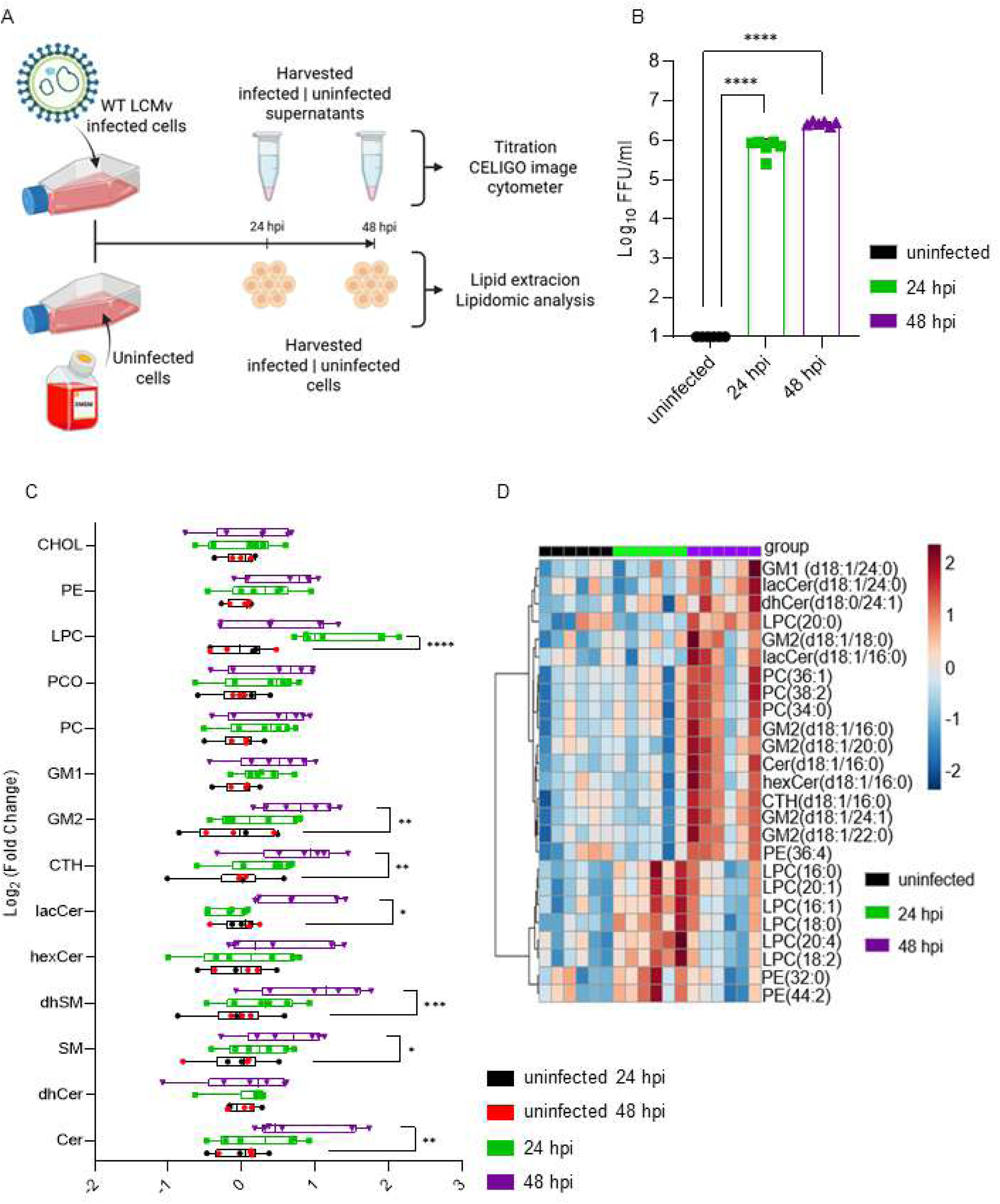
Global lipidomic changes in LCMV-infected A549 cells. **A.** Experimental design and sample collection. A549 cells were infected with LCMV-ARM (MOI = 0.05) or mock infected (DMEM). Cell culture supernatants (CCS) and cell monolayers were collected at 24 and 48 hpi for virus titration and lipid extraction, respectively. Figure created with Biorender. **B.** Production of LCMV infectious progeny at 24 and 48 hpi. Titers of infectious virus in CCS were determined by focus-forming assay (FFA). **C.** Box-and-whisker plots displaying the lipid subclasses detected in LCMV-infected and mock-infected cells (n= 6 per condition) at 24 and 48 hpi. Statistical significance was determined using the Holm-Sidak multiple comparison test. *, P < 0.05; **, P < 0.05; ***, P < 0.001. **D.** Heatmap displaying the top 25 lipid species selected by hierarchical clustering at 24 and 48 hpi. Heatmap colors represent relative lipid abundance, and values correspond to normalized, log2-transformed fold changes subjected to Pareto scaling. Each column denotes a single cell replicate (n = 6 for each group and time point).

### 3.3. Antiviral activity of lipid modulators against LCMV

Our transcriptomic (Fig.1F) and lipidomic (Fig. 3C, D) data revealed increased levels of several lipid species, particularly sphingolipids, following LCMV infection. To assess the functional relevance of these changes, we evaluated the effect on LCMV infection of pharmacological inhibitors of acetyl-CoA carboxylase (ACC) and neutral sphingomyelinase 2 (nSMase2), key enzymes in lipid synthesis and sphingolipid metabolism, respectively. For this, we determined the dose-dependent effect on LCMV multiplication of the ACC inhibitors PF-05175157 and CP-640186, which block fatty acid synthesis and impact multiple aspects of lipid biogenesis, and nSMase2 inhibitors DPTIP and cambinol, which inhibit the conversion of SM to Cer (Fig. 4A). All four inhibitors exhibited a dose-dependent inhibitory effect on LCMV multiplication (Fig. 4). Determination of EC_50_, and CC_50_ values for each inhibitor and their corresponding selectivity index (SI = CC_50_/EC_50_,), indicated that the antiviral activity of ACC (PF-0517515 and CP-640186) and nSMase2 (cambinol) inhibitors was not primarily caused by effects on cell viability.

**Figure 4.**
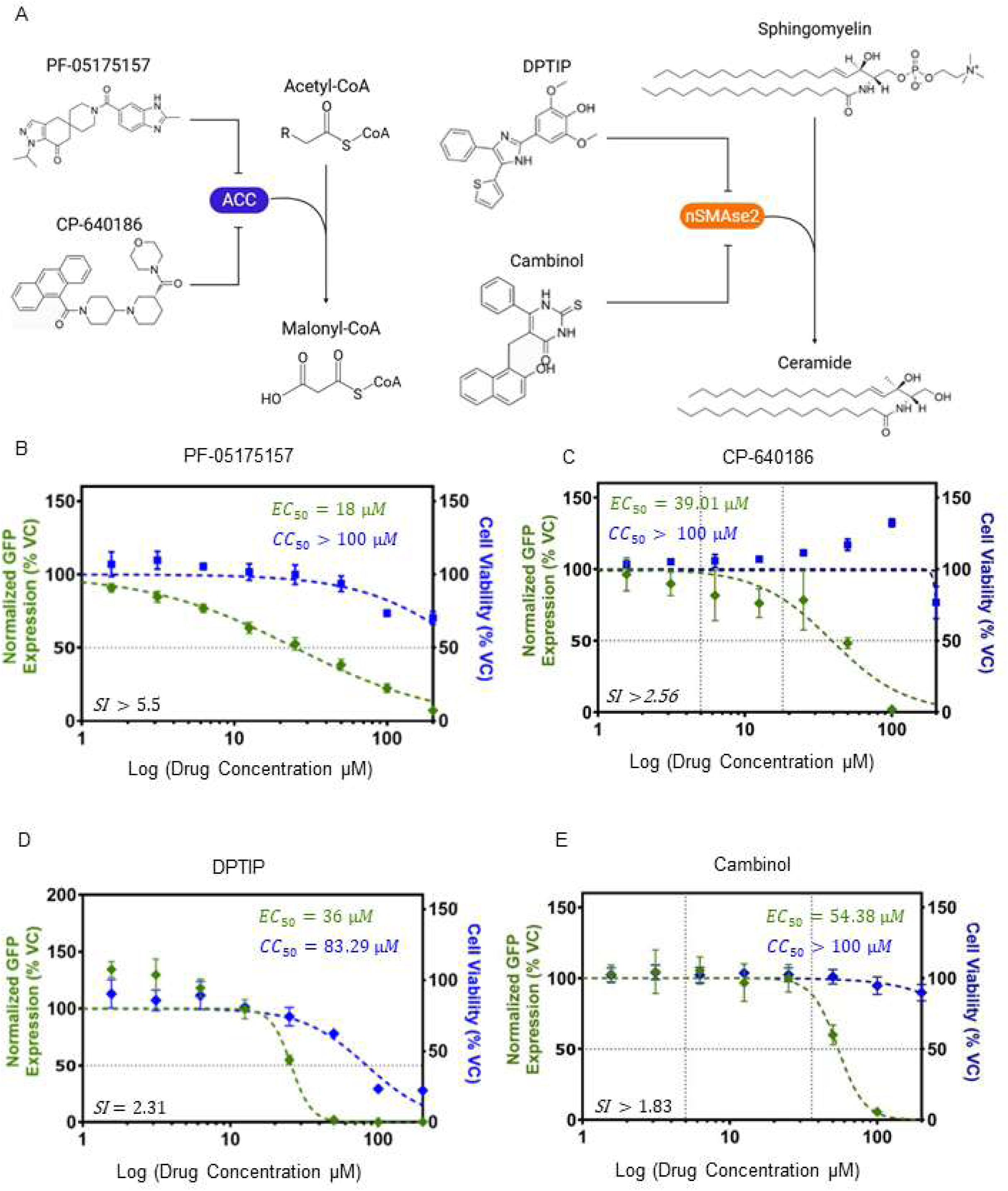
Antiviral activity of lipid modulators against LCMV. **A.** Schematic representation of the cellular pathways targeted by the tested inhibitors. **B-E.** Dose-dependent effect of the ACC inhibitors PF-05175157 (**B**) and CP-640186 (**C**), and nSMase2 inhibitors DPTIP (**D**) and Cambinol (**E**) on LCMV multiplication in Vero E6 cells. Cells were infected with rLCMV/GFP (MOI = 0.05) and treated with the indicated concentrations of each inhibitor. At 48 hpi, cell monolayers were fixed (PFA 4%), and numbers of infected cells were determined by GFP fluorescence. Cell viability was determined using an MTS/formazan bioreduction assay (n=3).

### 3.4. Chemical transcriptomics in LCMV-infected cells treated with lipid modulators

We next examined the effect of lipid modulators treatment on the transcriptomic profile of LCMV-infected cells. For this we selected the ACC (PF-05175157) and the nSMase2 (DPTIP) inhibitors as representative compounds affecting lipid metabolism via distinct mechanism of action (Fig. 5A). The concentration of PF-05175157 (18 µM) and DPTIP (36 µM) were selected based on their EC_50_ and CC_50_ values (Fig. 4B, 4D) and corresponded to concentrations at which the compounds exerted antiviral activity while having a negligible impact on cell viability. Treatment with the PF-05175157 and DPTIP altered the transcriptional profiles of LCMV-infected cells (Fig. 5B). Transcriptional shifts observed in infected cells treated with PF-05175157 or DPTIP were clearly distinct from the transcriptional profile observed in LCMV infected and vehicle control (VC) treated cells. We observed a reduction in the expression of genes related to GO:BP terms associated to response to stress, immune responses and defense responses (Fig. 5C), which was consistent with a reduction in viral load caused by the drug treatment. Commonly altered gene signatures (Fig. 5D) included a decrease in the expression of transcription factors related to histone deacetylase 1 activity (HDAC1, HDAC4), interferon signaling and regulation (STAT1, IRF1, STAT2), transcriptional regulation (EGR1, HIVEP2, CEBPD, MYC, E2F1, MYCN, JUN, SP1), regulation of lipid metabolism (PPARA), cellular response to interleukins (STAT3), endoplasmic reticulum stress (ATF4) or cellular stimuli such inflammation or immunity (RELA, NFKB1).

**Figure 5.**
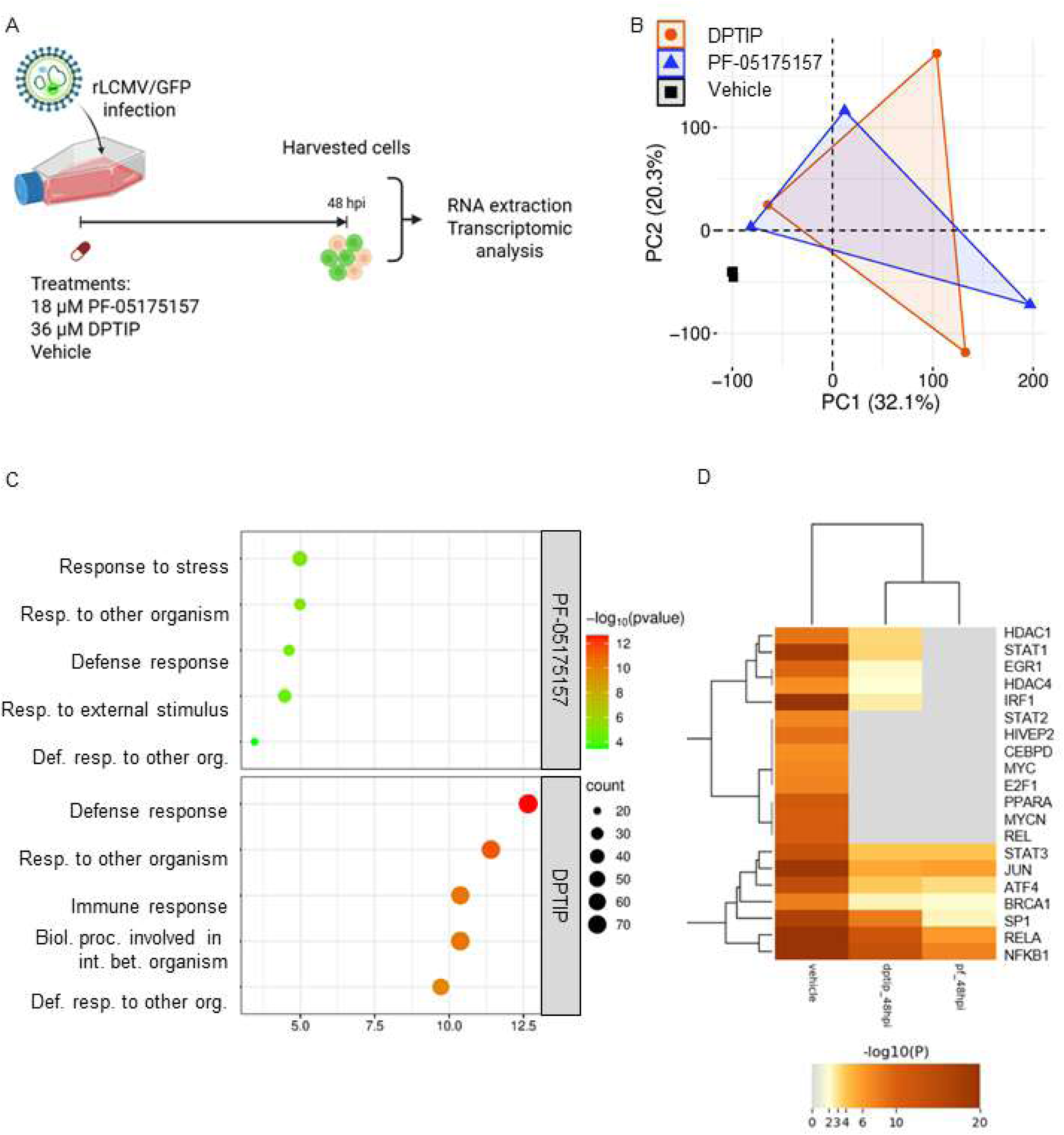
Chemical transcriptomic analysis of LCMV-infected cells treated with lipid metabolism modulators. **A.** Experimental design and sample collection. A549 cells were infected with rLCMV/GFP (MOI = 0.05) and treated with PF-05175157 (18 μM), DPTIP (36 μM) or vehicle control (VC). At 48 hpi total cellular RNA was isolated and analyzed by RNAseq. Figure created with Biorender. **B.** Principal component analysis (PCA) plot showing global gene expression profiles in infected-cells treated with the indicated inhibitors or VC **C.** Functional enrichment analysis of top 5 GO: BP terms corresponding to down-regulated DEGs in PF-05175157-and DPTIP-treated cells **D.** Heatmap showing enriched transcription factors for all the DEGs identified in inhibitor-and VC-treated cells using TRRUST database.

### 3.5. Effect of pharmacological inhibition of ACC and nSMase2 on LCMV multi-step growth kinetics

We next examined the effect of ACC (CP-640186) and nsMase2 (cambinol) inhibitors on LCMV multi-step growth kinetics. For this, we infected A549 cells with rLCMV/GFP-P2A-NP (MOI = 0.05) and treated them with different (25, 50 and 100 µM) concentrations of CP-640186 or cambinol, or with VC. We also used VC containing 1% DMSO to match the DMSO concentration present in the highest concentration of CP-640186 and cambinol. At the indicated hpi, we collected CCS and fixed cells with 4% PFA and stained them with DAPI. GFP expression levels and DAPI staining signals were quantified using a Cytation 5 plate reader (BioTek, Agilent), and their values were normalized to those of VC-treated and infected cells. Normalized GFP and DAPI values were used to determine virus infectivity and cell viability, respectively (Figs 6A and 7A). Mean ± SD values (*n* = 37 points) of GFP and DAPI signals are shown. We used CCS samples to determine virus titers (Figs. 6B and 7B). Treatment with cambinol (100 µM) reduced production of infectious viral progeny by ∼ 2 logs at 48 hpi and by 1 log at 72 hpi (Fig. 6B), which correlated with restricted virus propagation within the infected cell monolayer (Fig. 6 Ci) as determined by inhibition of GFP signal. Compared to VC treated cells, cambinol treatment reduced DAPI signal by ∼ 10, 30 and 40% in A549 cells at 24, 48 and 72 hpi (Fig. 6 Cii).Treatment with CP-640186 (100 µM) reduced production of infectious viral progeny by 1 log at 48 hpi (Fig. 7B), which correlated with restricted virus propagation within the infected cell monolayer (Fig. 7 Ci) as determined by GFP signal. Compared to VC treated cells, CP-640186 (100 µM) reduced DAPI signal by < 10% at all tested hpi (Fig. 7 Cii), indicating that CP-640186 anti-LCMV activity was not the result of compound associated cytotoxicity.

**Figure 6.**
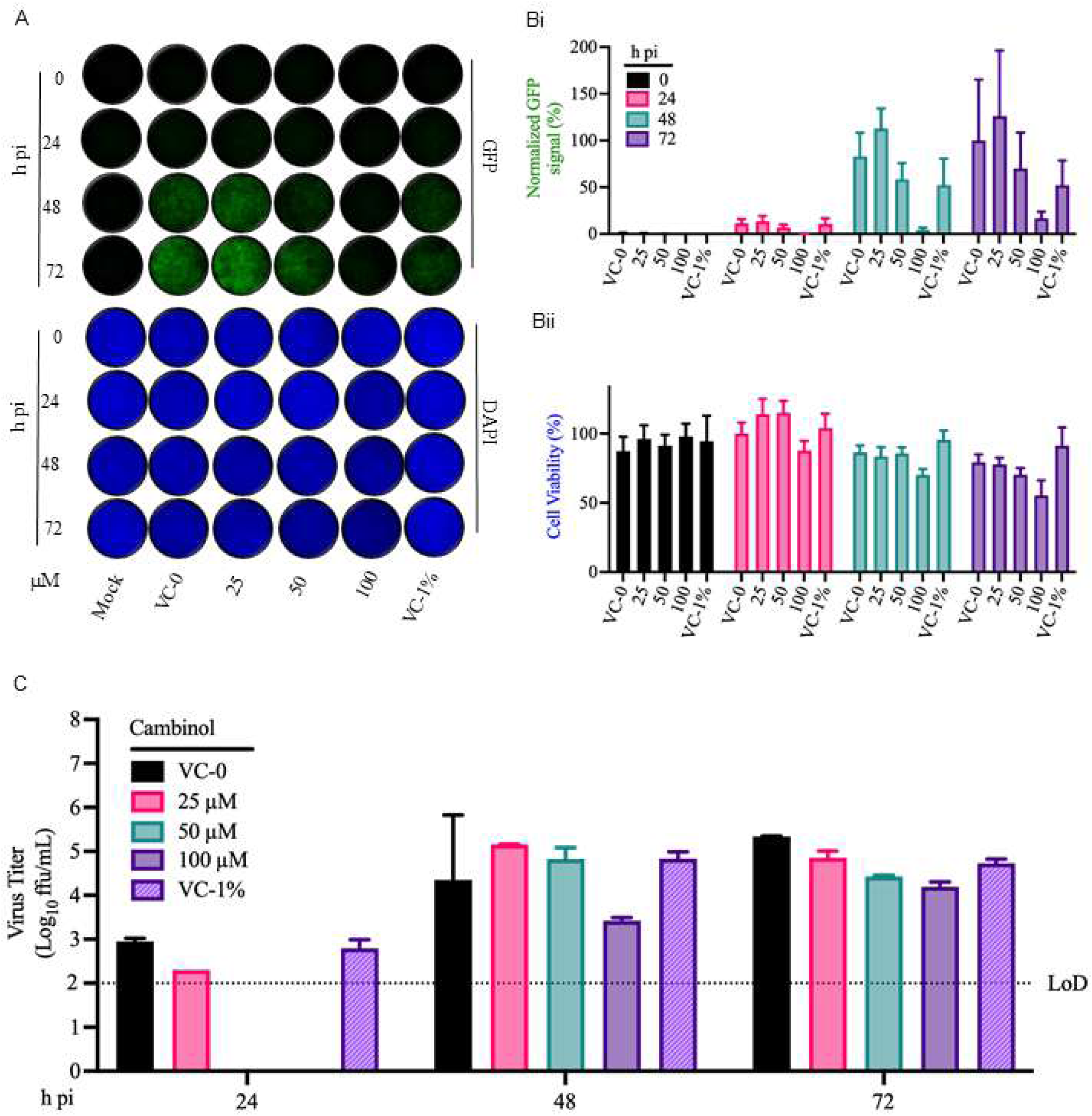
Effect of nSMase2 inhibition of LCMV multi-step growth kinetic in A549 cells. **A**. Effect of cambinol on production of infectious viral progeny. A549 cells were seeded at 5 × 10^5^ cells/well in an M12-well plate, infected (MOI = 0.05) with rLCMV/GFP-P2A-NP and treated with cambinol (25, 50 and 100 µM) or with VC or VC containing 1% DMSO. At the indicated time points, CCSs were collected, and titers of infectious virus were determined by FFA using Vero E6 cells. **B**. At the indicated hpi, samples from (**A**) were fixed with 4% PFA and stained with DAPI. Images (4× magnification) were acquired using a Keyence BZ-X710 microscope. **C**. GFP (**C**(**i**)) and DAPI (**C**(**ii**)) signals were quantified using Cytation 5 and normalized to the average value of the VC-treated samples at 72 hpi.

**Figure 7.**
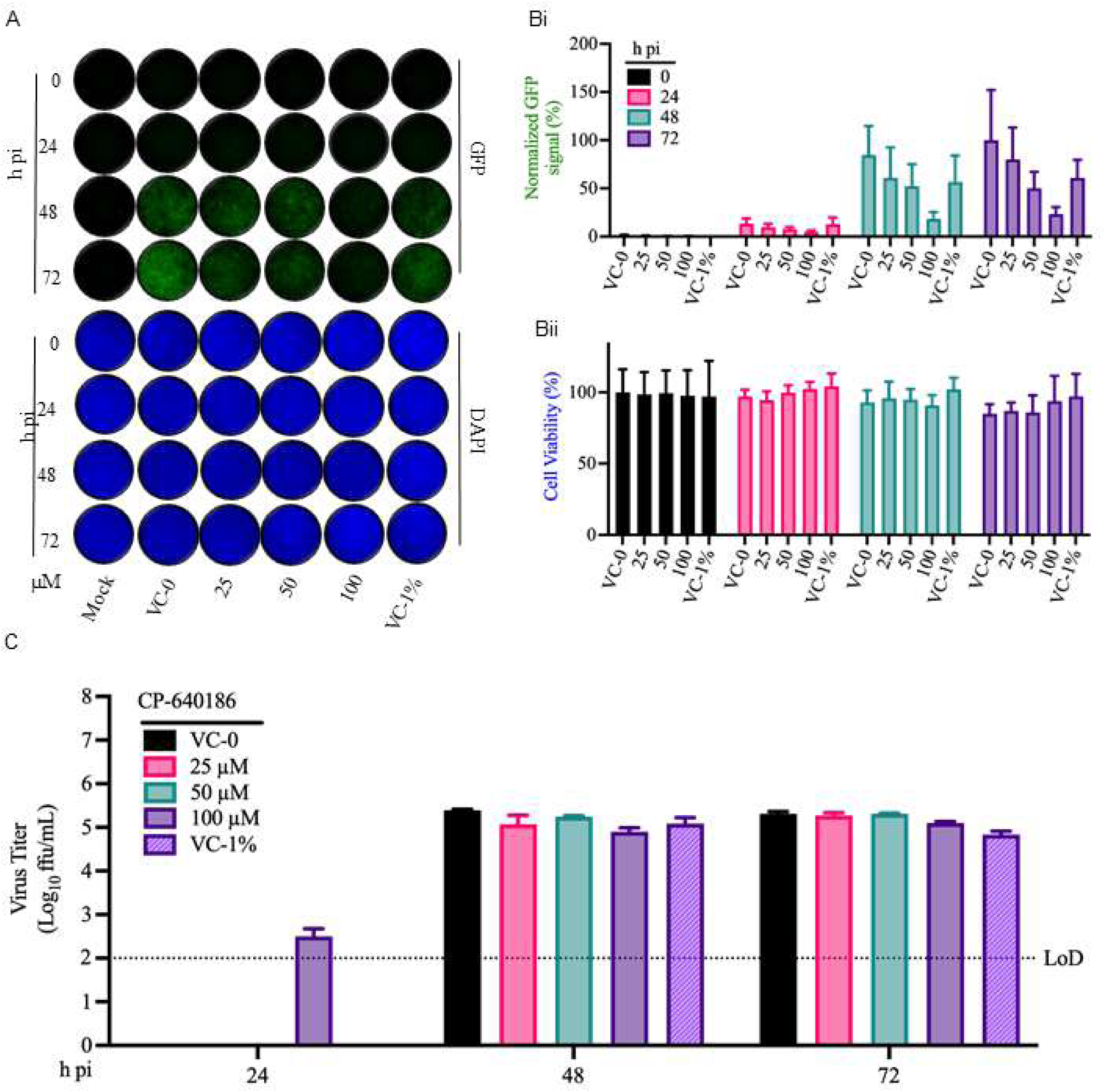
Effect of ACC inhibition of LCMV multi-step growth kinetic in A549 cells. **A**. Effect of CP-640186 on production of infectious viral progeny. A549 cells were seeded at 5 × 10^5^ cells/well in an M12-well plate, infected (MOI = 0.05) with rLCMV/GFP-P2A-NP and treated with CP-640186 (25, 50 and 100 µM) or with VC or VC containing 1% DMSO. At the indicated time points, CCSs were collected, and titers of infectious virus were determined by FFA using Vero E6 cells. **B**. At the indicated hpi, samples from (**A**) were fixed with 4% PFA and stained with DAPI. Images (4× magnification) were acquired using a Keyence BZ-X710 microscope. (**C**) GFP (**C**(**i**)) and DAPI (**C**(**ii**)) signals were quantified using Cytation 5 and normalized to the average value of the VC-treated samples at 72 hpi.

## 4. DISCUSSION

The role of lipids in replication of positive sense RNA virus has been extensively documented (Stancheva & Sanyal, 2024). In contrast, there is limited information regarding the interactions between cellular lipids and negative strand RNA viruses (Tam et al., 2013; Zhao et al., 2026). In this study, we combined transcriptomics, lipidomics, and pharmacological perturbation to investigate the interaction between the prototypic MaAv LCMV and the host cell lipid metabolism. During the preparation of this manuscript, a lipidomic analysis of Vero cells infected with a r3LCMV LASV chimera was reported, showing alterations in lipid composition at a single infection time point (72 h pi) (Noble et al., 2026). In contrast, our results analyzing different times post-infection revealed a temporal pattern of lipid changes throughout infection. Early stages of infection were characterized by increased levels of lysophospholipids, including single-acyl glycerophospholipids (LPLs) and lysophosphatidylcholine (LPC), whereas at later stages of infection were associated with accumulation of multiple sphingolipids species, including Cer, dhCer, SM, lacCer, gangliosides (GM1/GM2), and hexCer. These findings suggest that LCMV infection induces a temporally regulated remodeling of membrane lipid composition. Importantly, our results demonstrate for the first time that LCMV infection profoundly reshapes the cellular lipid landscape, affecting not only fatty acid synthesis but also the sphingolipid metabolic pathways. Altered levels of sphingolipids and LPCs have been reported during infection with other viruses (Requena et al., 2025), and sphingomyelin synthase 1 (SGMS1) has been identified as a proviral factor in LCMV-infected cells (Han, S. et al., 2024). This dependence on sphingolipids is consistent with the established importance of membrane lipid microdomains, enriched in cholesterol and sphingolipids (Gee et al., 2023), which have been involved in MaAv assembly (Agnihothram et al., 2009; Cordo et al., 2013)

Pharmacological targeting of lipid synthesis using ACC inhibitors, which block the rate-limiting step of fatty acid synthesis, demonstrated that LCMV infection dependents on *de novo* fatty acid biogenesis, a finding consistent with the requirement of specific lipid species to support efficient LCMV infection (Noble et al., 2026). Fatty acids provide the building blocks of more complex lipids such as sphingolipids and glycerophospholipids. In LCMV-infected mice, serum metabolomics revealed marked perturbations in circulating glycerophospholipids and sphingolipids during the peak of hepatitis, accompanied by elevated cholesterol levels (Ripple et al., 2017). These infection-induced changes in complex lipids engaged the lipid sensor TREM2 on myeloid cells, thereby exacerbating liver immunopathology and indirectly influencing viral control despite comparable T cell responses (Ripple et al., 2017). These observations suggest that virus-induced remodeling of sphingolipid and glycerophospholipid pools links fatty acid metabolism to a host lipid environment that modulates LCMV-associated disease outcomes (Ripple et al., 2017).

Although with different potencies, all tested lipid synthesis inhibitors (PF-05175157, CP-640186, DPTIP and cambinol) restricted LCMV infection in a dose-dependent manner. Treatment with 100 µM cambinol reduced production of infectious viral progeny by 2 logs at 48 hpi, an observation that cannot be accounted by cytotoxicity as no significant effect on cell viability was observed under these conditions (Fig. 4E). Altered levels of sphingolipids and LPCs have been reported during infection with other viruses (Requena et al., 2025), and sphingomyelin synthase 1 (SGMS1) has been identified as a proviral factor in LCMV-infected cells (Han, S. et al., 2024). Consistent with their known roles in membrane organization, sphingolipids enrichment in LCMV-infected cells may facilitate the formation of membrane microdomains enriched in cholesterol and sphingolipids (Gee et al., 2023), which can support assembly of viral replication complexes and budding. In our cell-based infection assay cambinol has an EC_50_ of ∼ 54 µM, whereas in a biochemical assay cambinol was reported to have an IC_50_ of 5 µM for nSMase2 (Stepanek et al., 2019). These apparent conflicting findings likely reflect differences in assay conditions.

Overall, our findings revealed that LCMV induces a dynamic rewiring of lipid metabolic processes, prominently affecting sphingolipid metabolism, and that pharmacological targeting of these pathways restricts infection in cultured cells, providing proof-of-concept that lipid pathways are druggable targets for the potential development of HDAs to treat infections by human pathogenic MaAv. targets. However, the relatively high concentrations of ACC and nSMase2 inhibitors required for their observed anti-LCMV activity may limit their immediate translational application without structure-activity optimization efforts and validation studies in primary cells and animal models of infection.

## Acknowledgements

Not applicable

## Author contributions

**Patricia Mingo-Casas:** Conceptualization, Formal analysis, Funding acquisition, Investigation, Writing – original draft, Writing – review and editing. **Haydar Witwit:** Formal analysis, Investigation, Writing – review and editing. **Mireia Casasampere**: Formal analysis, Investigation, Writing – review and editing. **Ana-Belén Blázquez:** Formal analysis, Project administration, Supervision, Writing – review and editing. **Beatrice Cubitt:** Investigation, Writing – review and editing. **Miguel A. Martín-Acebes:** Conceptualization, Formal analysis, Funding acquisition, Supervision, Writing – original draft, Writing – review and editing. **Juan Carlos de la Torre:** Conceptualization, Formal analysis, Supervision, Writing – original draft, Writing – review and editing.

## Data availability

RNA-seq datasets generated in this work are available at GEO (https://www.ncbi.nlm.nih.gov/geo/) under Accession GSE334699.

## Figure legends

**Figure S1. Changes in the lipidome of A549 cells infected with LCMV (MOI = 0.05) at 24 and 48 hpi or mock-infected.** Lipids were identified and quantified by mass spectrometry as described in Materials and Methods. The heatmap was created using Metaboanalyst 6.0. Replicates for each condition are displayed (n = 6).

