## Supplemental Figure 1 for "Mammarenavirus-Induced Remodeling of the Cellular Lipid Landscape Reveals Sphingolipid Metabolism as a Novel Target for Antiviral Intervention"

**Supplementary figure legend**

**Figure S1. Heatmap of the lipidome of Vero cells infected with LCMV**. Lipid levels in the scales denote normalized, $log_2$-transformed fold change and Pareto-scaled values for uninfected cells, and LCMV-infected cells at 24 and 48 hpi. Columns denote analyzed samples and rows lipid species ordered by subclass and total carbon. Samples were grouped by hierachical clustering.

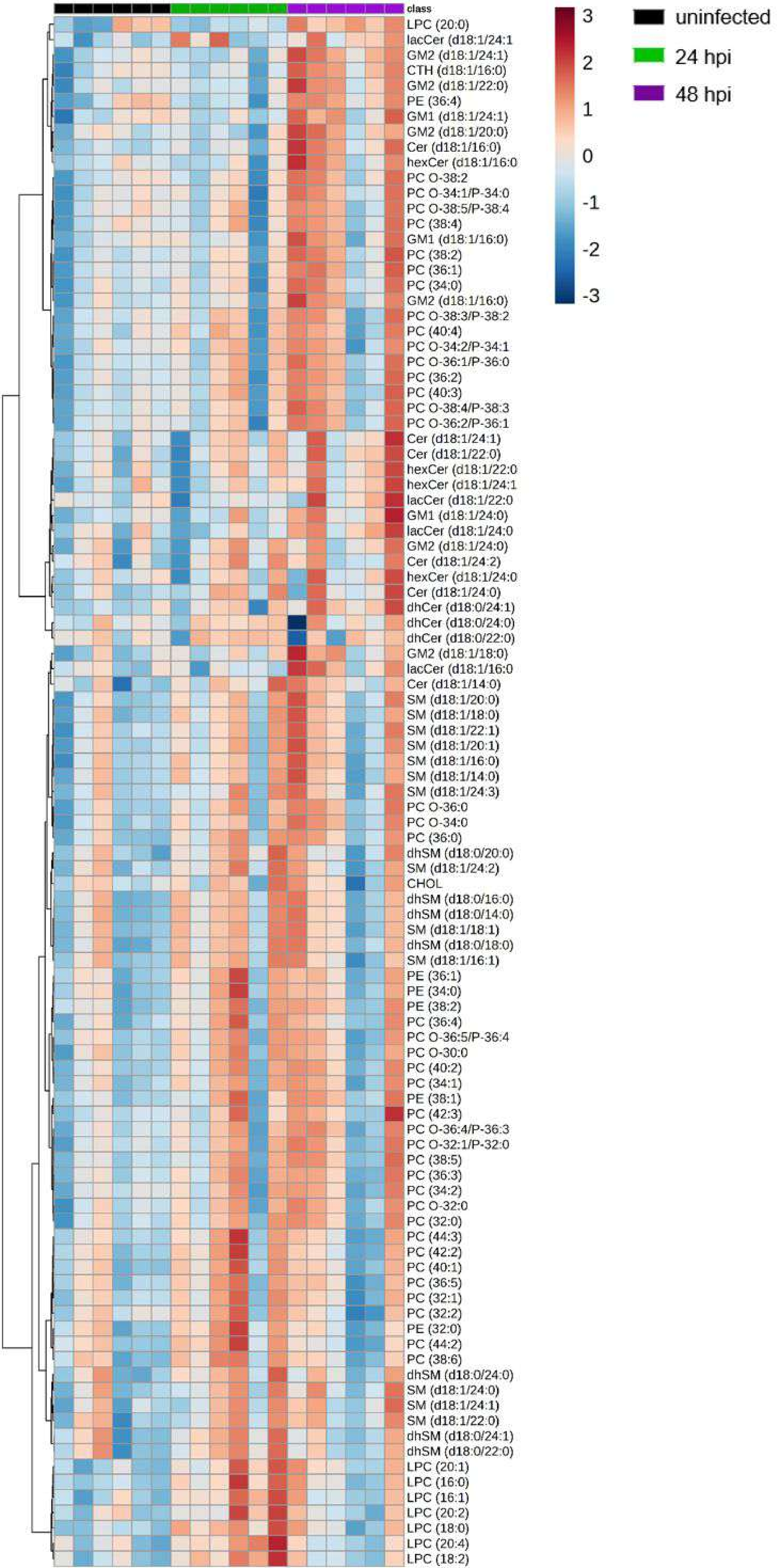
class
uninfected
24 hpi
48 hpi
3
2
1
0
-1
-2
-3
LPC (20:0)
lacCer (d18:1/24:1
GM2 (d18:1/24:1)
CTH (d18:1/16:0)
GM2 (d18:1/22:0)
PE (36:4)
GM1 (d18:1/24:1)
GM2 (d18:1/20:0)
Cer (d18:1/16:0)
hexCer (d18:1/16:0
PC O-38:2
PC O-34:1/P-34:0
PC O-38:5/P-38:4
PC (38:4)
GM1 (d18:1/16:0)
PC (38:2)
PC (36:1)
PC (34:0)
GM2 (d18:1/16:0)
PC O-38:3/P-38:2
PC (40:4)
PC O-34:2/P-34:1
PC O-36:1/P-36:0
PC (36:2)
PC (40:3)
PC O-38:4/P-38:3
PC O-36:2/P-36:1
Cer (d18:1/24:1)
Cer (d18:1/22:0)
hexCer (d18:1/22:0
hexCer (d18:1/24:1
lacCer (d18:1/22:0
GM1 (d18:1/24:0)
lacCer (d18:1/24:0
GM2 (d18:1/24:0)
Cer (d18:1/24:2)
hexCer (d18:1/24:0
Cer (d18:1/24:0)
dhCer (d18:0/24:1)
dhCer (d18:0/24:0)
dhCer (d18:0/22:0)
GM2 (d18:1/18:0)
lacCer (d18:1/16:0
Cer (d18:1/14:0)
SM (d18:1/20:0)
SM (d18:1/18:0)
SM (d18:1/22:1)
SM (d18:1/20:1)
SM (d18:1/16:0)
SM (d18:1/14:0)
SM (d18:1/24:3)
PC O-36:0
PC O-34:0
PC (36:0)
dhSM (d18:0/20:0)
SM (d18:1/24:2)
CHOL
dhSM (d18:0/16:0)
dhSM (d18:0/14:0)
SM (d18:1/18:1)
dhSM (d18:0/18:0)
SM (d18:1/16:1)
PE (36:1)
PE (34:0)
PE (38:2)
PC (36:4)
PC O-36:5/P-36:4
PC O-30:0
PC (40:2)
PC (34:1)
PE (38:1)
PC (42:3)
PC O-36:4/P-36:3
PC O-32:1/P-32:0
PC (38:5)
PC (36:3)
PC (34:2)
PC O-32:0
PC (32:0)
PC (44:3)
PC (42:2)
PC (40:1)
PC (36:5)
PC (32:1)
PC (32:2)
PE (32:0)
PC (44:2)
PC (38:6)
dhSM (d18:0/24:0)
SM (d18:1/24:0)
SM (d18:1/24:1)
SM (d18:1/22:0)
dhSM (d18:0/24:1)
dhSM (d18:0/22:0)
LPC (20:1)
LPC (16:0)
LPC (16:1)
LPC (20:2)
LPC (18:0)
LPC (20:4)
LPC (18:2)
